# Cell type matching in single-cell RNA-sequencing data using FR-Match

**DOI:** 10.1101/2021.10.17.464718

**Authors:** Yun Zhang, Brian Aevermann, Rohan Gala, Richard H. Scheuermann

## Abstract

Reference cell type atlases powered by single cell transcriptomic profiling technologies have become available to study cellular diversity at a granular level. We present FR-Match for matching query datasets to reference atlases with robust and accurate performance for identifying common and novel cell types and suboptimally clustered cell types in the query data. FR-Match shows excellent performance for cross-platform, cross-sample type, and cross-tissue region cell type matching.

## Main

Single cell transcriptomic profiling has emerged as a powerful tool to characterize the cellular heterogeneity in complex biological systems. Large collaborative consortia, including the Human Cell Atlas [1] and NIH BRIAN Initiative [2, 3], have adopted the unbiased single-cell/nucleus RNA-sequencing (sc/snRNA-seq) technologies to generate reference cell type atlases at single cell resolution across many organs and species at an unprecedented level of granularity; a series of recent publications have reported 128 transcriptomically-distinct cell types in human primary motor cortex (M1) [4], 116 cell types in mouse primary motor cortex (MOp) [5], and 75 cell types in human middle temporal gyrus (MTG) neocortex [6]. The Allen Institute for Brain Science has made these comprehensive datasets available to serve as reference cell type atlases (https://portal.brain-map.org/atlases-and-data/rnaseq).

An important role for these reference datasets is to support the matching of new query data to the reference cell types using recently-developed computational methods. Azimuth is a web application for reference-based single-cell analysis following the Seurat pipeline [7]. Online iMNF is an extension of the Liger pipeline for single-cell multi-omics integration using iterative online learning [8, 9]. ScArches is a deep learning strategy for mapping query datasets on top of a reference by single-cell architectural surgery [10]. The mathematical foundation of these methods are linear algebra techniques (canonical correlation analysis (CCA) for Seurat and non-negative matrix factorization (NMF) for Liger) that effectively decompose the structure of large data matrices for integrative analysis. While these methods are great tools for single-cell data integration, producing integrated UMAP visualization for both query and reference datasets with minimal batch effects, cell type matching is a more pragmatic use case that requires not only integrating the query cells into the reference, but also being able to make a clear distinction between common and novel cell types existing in the query dataset and the studied conditions.

Previously, we reported a computational pipeline for downstream cell type analysis of scRNA-seq data combining NS-Forest, a random forest machine learning algorithm for the identification of minimum sets of marker genes for given cell types [11, 12], and FR-Match, a minimum spanning tree-based statistical testing approach for cell type matching of query and reference datasets, and demonstrated its performance in matching cell types from overlapping brain regions [13]. We introduced the concept of cell type “barcodes” [12, 13] using NS-Forest marker genes to visualize and characterize the distinction between different cell types [12]. The NS-Forest marker genes also serve as a feature selection approach for FR-Match that essentially matches the query and reference cell types based on the cell type barcode gene expression patterns [13]. Here, we report recent enhancements to FR-Match, including a normalization step and a cell-to-cluster matching scheme, and show that the cell type barcodes provide evidence and explainability for the matching results. The enhanced FR-Match was found to effectively match cell types between platforms (10X and SMART-seq; scRNA-seq and smFISH), sample types (whole cells and nuclei), and tissue regions (human M1 and MTG) and provided evidence for suboptimal partitioning in the clustering step.

We designed a normalization procedure based on the marker gene expression patterns observed in the cell type barcode plots, to dampen technical artifacts observed in different scRNA-seq platforms. We observed that barcode plots from the SMART-seq platform (Figure 1A(i)) and the 10X platform (Figure 1A(iv)) showed similar marker gene expression specificity, but different expression distributions (Supplemental Figure 1) and variable non-specific background expression. To address these technical artifacts for cell type matching, min-max rescaling is applied to each gene independently for both SMART-seq and 10X data, to globally align the data in the range of [0, 1]. The SMART-seq platform showed better sensitivity for low expression genes than the 10X experiment, but also showed more background noise. To reduce the background noise while preserving the expression signals, the normalization step for the SMART-seq data uses a per-barcode per-gene summary statistic (mean or median) to weight the barcode pattern by multiplying the summary statistic to the gene expression submatrix of that cluster (see Methods). In the case of median, the result if that for genes expressed in a minority of cells (median = 0), expression in these cells is set to zero. Finally, the weighted barcode is again rescaled to [0, 1] for matching. The above procedure effectively aligned the cross-platform barcode patterns (Figure 1A(ii)(iii)), producing similar signal and noise levels.

**Figure 1:**
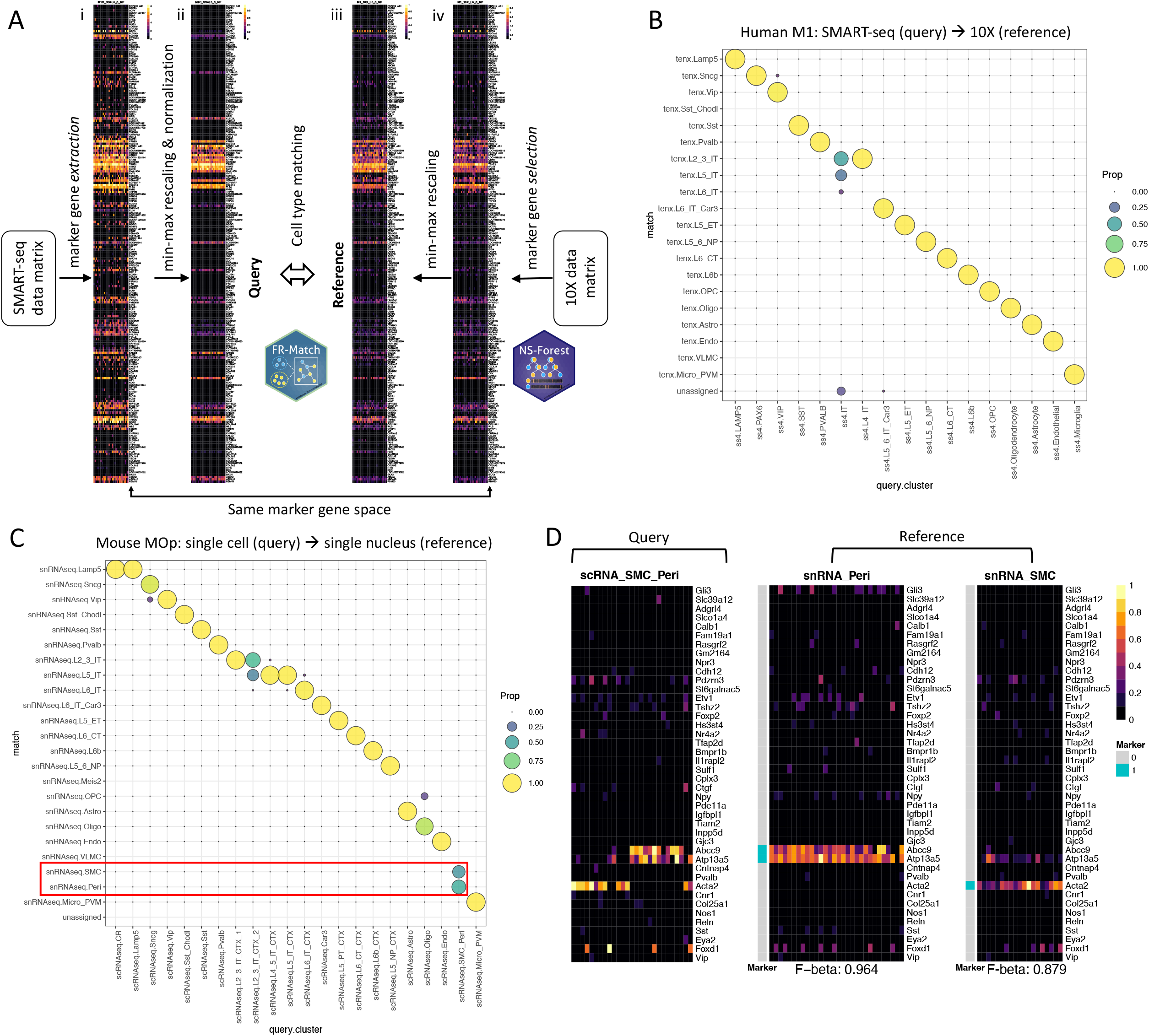
Cell type matching based on features selected using NS-Forest and feature distribution matching using FR-Match. **A.** Schematic of the cross-platform cell type matching pipeline. The pipeline includes input reference data from the 10X Genomics platform, marker gene selection using NS-Forest algorithm based on the reference gene expression data, marker gene expression extraction for the query input SMART-seq platform data using the 10X reference marker genes, platform-specific rescaling and normalization steps, and marker gene expression distribution matching of the query and reference normalized data using the FR-Match algorithm. **B.** Cell-to-cluster matching results for matching cell types from SMART-seq (query) to 10X (reference) datasets for the human M1 brain region using FR-Match. Results are shown as the proportion of cells matched between pairs of query and reference subclass cell types. Most of the query cells are matched with the expected reference cell type subclass, aligning diagonally in the plot. The only exception is the agglomerated query IT subclass that was matched to several layer-specific reference IT subclasses or unassigned. **C.** Cell-to-cluster matching results for matching cell types from single-cell RNA-seq (scRNA-seq) (query) to single-nucleus RNA-seq (snRNA-seq) (reference) 10X datasets for mouse MOp brain region using FR-Match. Highlighted in the red box is an example of evidence for an under-partitioned query SMC-Peri subclass in which cells were matched to either the SMC subclass or the Peri subclass in the reference dataset. **D.** Cell type barcodes of the query SMC-Peri subclass and the corresponding reference SMC subclass and reference Peri subclass. The barcode plots clearly show two distinct expression patterns in the query cluster, each reflecting one of the two reference cluster expression pattern, supporting the under-partitioning hypothesis.

To augment the original cluster-to-cluster matching in FR-Match, a cell-to-cluster extension was added based on an iterative procedure that allows each cell in the query cluster to be assigned a summary p-value, quantifying the confidence of matching to a reference cluster (see Methods). This extension is available as a stand-alone function “FRmatch_cell2cluster()” in the “FRmatch” R package (https://github.com/JCVenterInstitute/FRmatch). A cosine distance option was also added for robust matching between experiments with global data variabilities.

Using these pipelines and extensions, the cross-platform matching approach was validated using Allen human M1 snRNA-seq data generated using the *10X Chromium v3* protocol [4] as the reference and an M1 snRNA-seq dataset from another Allen study on multiple human cortical regions using the *SMART-seq v4* protocol [6] as a query. Although the raw counts of the query and the reference data showed very different data distributions (Supplementary Figure 1) the FR-Match matching results produced almost all one-to-one matching at the subclass level for all query cells (Figure 1B), with the exception of the agglomerated IT query type. Due to the grouping (under-partitioning) of the layer-non-specific IT cells in the query, the majority of these cells were matched to one of the different layer-specific IT reference types. These results verify that the normalization step for aligning SMART-seq and 10X data is effective and the extended FR-Match is robust to perform cross-platform cell type matching.

We also applied this FR-Match pipeline to assess cross-sample type matching using an *snRNA-seq* dataset from the Allen mouse MOp [5] as reference cell types and an *scRNA-seq* dataset from the MOp subset of a cell type taxonomy of the entire adult mouse isocortex and hippocampus [14] as the query. Since both datasets were generated using the 10X protocol, we only applied the min-max scaling in the normalization step. For subclass types, most of the query types were one-to-one matched to a reference type (Figure 1C). The highlighted box shows that query SMC-Peri cells were matched to either the SMC or Peri types in the reference, with ~50:50 split. A one-to-many match may indicate that the query cluster is under-partitioned [13]. An examination of the cell type barcode plots for these query and reference cell types (Figure 1D) showed two distinct patterns in the query barcode, each corresponding to one of the two reference barcodes supporting the under-partitioning hypothesis. Thus, the FR-Match cell type matching pipeline, together with the cell type barcode, showed excellent matching of single nucleus and single cell clusters and provided solid evidence of suboptimal partitioning based on marker gene expression in the matching results.

We benchmarked the FR-Match pipeline with the Azimuth and Online iNMF approaches for identification of suboptimal parititoning using human M1 (Supplementary Figure 2A-B) and mouse MOp (Supplementary Figure 2C-D) datasets. All cells were matched, but these integration methods were not able to split the under-partitioned clusters. In the mouse MOp case, Azimuth matched all query SMC-Peri cells to the reference Peri subclass with a few mismatched to the reference VLMC subclass (Supplementary Figure 2C). The Online iNMF produced joint clustering of the integrated data instead of explicitly reporting the cell-to-cell mapping. All the query SMC-Peri cells and the reference SMC and Peri cells were grouped in the same cluster from the joint clustering (Supplementary Figure 2D).

For the above two use cases, we also matched the most granular cell types and benchmarked in comparison with Azimuth. In the mouse MOp use case, the FR-Match results formed a clean diagonal alignment of cell types and assigned unmatched cells as “unassigned” in the bottom row (Figure 2A); in some cases with no matches to the reference clusters suggesting the presence of novel cell types in the query data. The Azimuth results also showed the majority of the matching cells along the diagonal, but with many more suboptimal matches scattered off-diagonal and no indication of novel unassigned cell types in the query data (Figure 2B). Similar results for human M1 can be found in Supplementary Figure 3.

**Figure 2:**
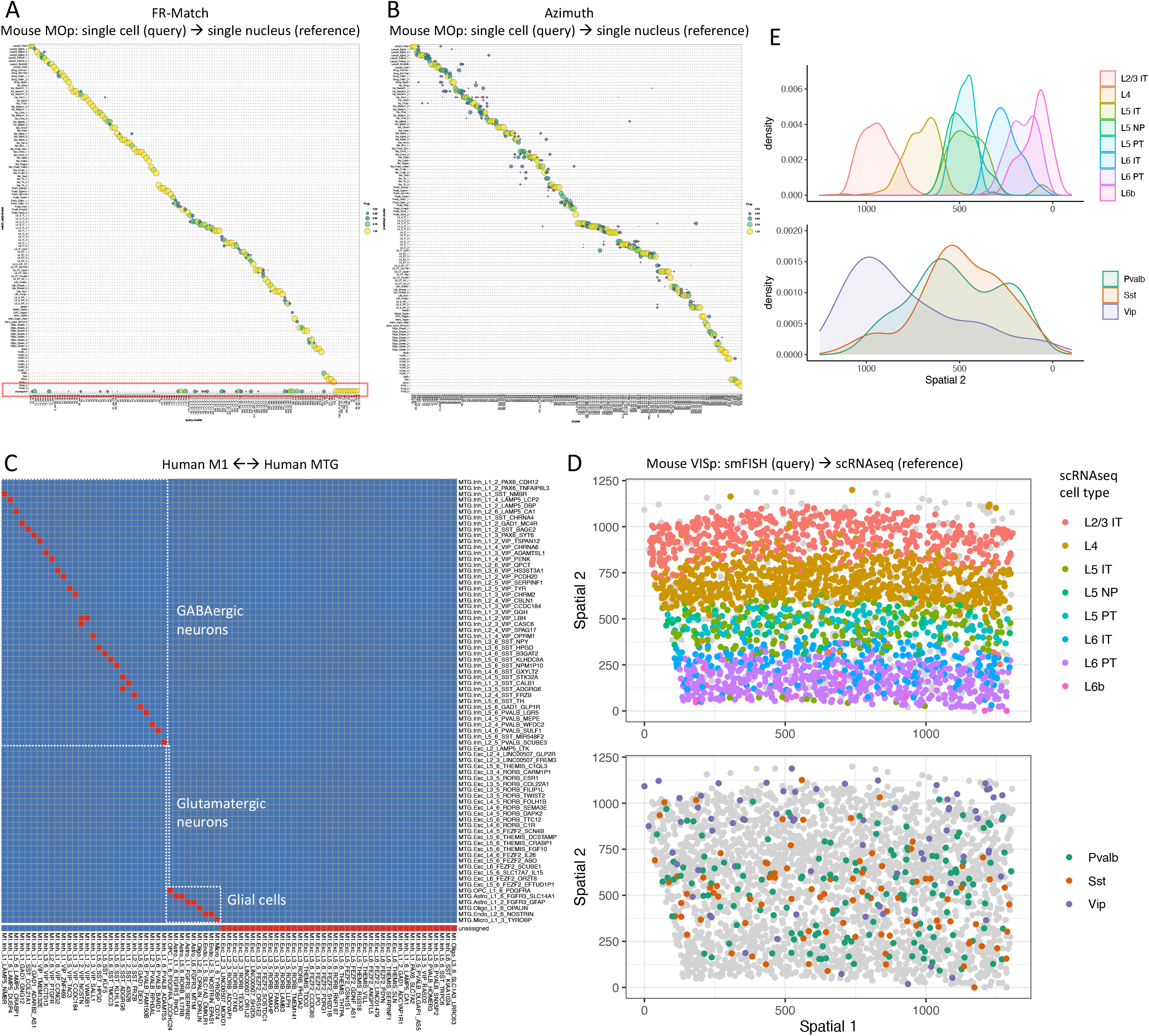
Additional FR-Match use cases and comparisons. **A.** Cell-to-cluster matching results for matching cell types from scRNA-seq (query) to snRNA-seq (reference) datasets of mouse MOp cell types at the most granular cell type resolution using FR-Match. The majority of cells in the query cell types were matched uniquely to reference cell types, showing clean diagonal matches with few off-diagonal matches. The highlighted box (in red) at the bottom is the “unassigned” row for the query cells that were not matched to any of the reference cell types based on the FR-Match results. The unassigned cells may correspond to novel query cell types not presented in the reference cell types. **B.** Matching results for matching cell types from scRNA-seq (query) to snRNA-seq (reference) datasets of mouse MOp cell type matching at the most granular cell type resolution using Azimuth. Though the majority of the cells were matched along the diagonal, there were many off-diagonal matches suggesting lower-quality matching. **C.** Cluster-to-cluster two-way matching results for matching cell types across the human M1 and MTG brain regions using FR-Match. FR-Match results suggests that most of the GABAergic and glial cell types are conserved across brain regions, whereas the glutamatergic cell types appear to be region-specific. **D.** Application of FR-Match to the spatial transcriptomics data. Cell type assignment of a mouse VISp smFISH dataset using scRNA-seq-defined reference cell types of the same brain region and the FR-Match cell-to-cluster algorithm. The assigned excitatory cell types clearly recapitulate the laminar distribution in the spatial coordinates (top); the assigned inhibitory cell types show the expected scattered spatial distribution. **E.** Spatial distributions of the excitatory cell types (top) and inhibitory cell types (bottom) summarized from the FR-Match cell type assignment results for the smFISH data shown in D.

Another important matching challenge is to match cell types across different tissues or anatomic regions within a tissue. The cluster-to-cluster version of FR-Match [13] was used to match cell clusters from the SMART-seq platform between the human M1 [4] and MTG [6] brain regions and the bi-directional (M1 as query to MTG as reference, and vice versa) matching results are shown in Figure 2C. Most of the GABAergic neuron and all of the glial cell types were nicely matched across these two cortical brain regions, but none of the glutamatergic neuron types were matched. This suggests that the inhibitory neuron and glial cell types are conserved across brain regions, whereas the excitatory neurons are cortical region specific. We also examined the cell type barcode plots for pairs of matched cell types showing highly similar expression patterns of the matched types using reciprocal marker genes, even though the best marker gene sets selected for each brain regions may be different when defined based on the cell types present in the dataset (e.g., Supplementary Figure 4).

Finally, we report the application of FR-Match for matching spatial transcriptomics data generated by single molecular fluorescence in situ hybridization (smFISH) [15] to a SMART-seq scRNA-seq dataset as the reference, both from mouse primary visual cortex (VISp) [16]. *De novo* clustering of the smFISH data was used to obtain broadly-defined cell type clusters. The FR-Match cell-to-cluster pipeline was used to assign a reference cell type to each spatial cell in the initial clusters. The FR-Match results successfully recapitulated the clear laminar distributions of excitatory neurons, corresponding to their assigned cell types (Figure 2D-E). In contrast, the inhibitory neurons were scattered across all layers, with the *Vip* type located more densely in upper layers and the *Sst* and the *Pvalb* types located more densely in deeper layers. The FR-Match cell type assignment for the spatially sequenced cells reflected their expected laminar patterns.

In summary, we extended our cell type matching pipeline to perform both cell-to-cluster and cluster-to-cluster matching. The added normalization step and cosine distance option allow FR-Match to perform robust and accurate cell type matching across platforms (SMART-seq with 10X), sample types (single-cell with single-nucleus), brain regions (M1 with MTG), and data modalities (spatial transcriptomics with scRNA-seq). Compared with other methods, FR-Match can effectively detect non-optimally partitioned clusters from the previous clustering step, and uniquely identify potential novel cell types as “unassigned” cells. The cell type barcodes can be useful for investigators to interpret the underlying transcriptomic drivers of the FR-Match results, assisting future research directions.

## Methods

### FR-Match cell-to-cluster matching algorithm

As originally conceived, FR-Match is a cluster-to-cluster matching algorithm that utilizes a graphical model and minimum spanning trees to determine the data distributional equivalence between two cell type clusters derived from single cell or single nucleus RNA sequencing (scRNA-seq) data in multivariate space [13]. The required input data for FR-Match are cell-by-gene expression matrices and cell cluster membership labels for both query and reference data. The output of the original FR-Match is a map between the query cluster and the reference cluster labels, thus assigning known reference cell types to the query cell clusters, or defining a query cluster as an unassigned “novel” cell type not found in the reference.

Here, we extend the FR-Match algorithm to map each query *cell* to the known cell type clusters in the reference, i.e., cell-to-cluster matching. The input data are the same as before. If the query clusters are unavailable, it is sufficient to obtain broadly-grouped clusters by using the popular Louvain [17] or Leiden [18] clustering algorithms for scRNA-seq data. These clusters may not be at the ideal level of granularity to be directly matched to the granular cell types defined in the reference; rather, they provide candidate cluster memberships as the input data to FR-Match.

The extended cell-to-cluster FR-Match algorithm is implemented in the function `FRmatch_cell2cluster()`, and its plotting function implemented as `plot_FRmatch_cell2cluster()` in the `FRmatch R` package. The steps of the algorithm and the corresponding arguments in the functions are as follows:

1. Dimensionality reduction:
  1.1. Select informative marker genes using the companion marker gene selection algorithm - NS-Forest - or user-defined marker genes for the reference dataset;
  1.2. Extract the expression data for the reference marker genes in the query dataset, i.e., project the query data into the reference feature space for reduced dimensionality;
2. Pairwise iterative matching:
  2.1. For each pair of query cluster (*j*) and reference cluster (*k*):
    2.1.1.For subsample iteration index *i* iterating from 1 to the total number of iterations (`subsamp.iter=`):
      2.1.1.1. Subsample the same number of cells (`subsamp.size=`) from the query and reference clusters, denoted as *S*_*i*_ for the set of selected query cells;
      2.1.1.2. Perform Friedman-Rafsky test (FR test) [19], a nonparametric statistical test for multivariate two-group comparison, and obtain p-value from the test, denoted as *p*_*i*_;
      2.1.1.3. Assign the p-value to the selected query cells, i.e., *p*_*ck*_ = *p*_*i*_ for *c* ∈ *S*_*i*_ and reference cluster *k*;
      2.1.1.4. Repeat 2.1.1.1 and 2.1.1.2, and obtain *p*_*i′*_ for the updated iteration *i*′;
      2.1.1.5. Update pck *p*_*ck*_ = max {*p*_*ck*_ . *p*_*i′*_) for *c* ∈ *S*_*i′*_ sir and reference cluster *k*, i.e., re-assign *p*_*ck*_ if *p*_*i′*_ is greater than previously assigned *p*_*ck*_;
    2.1.2.End looping over iterations;
  2.2. End looping over query-and-reference-cluster-pairs;
  2.3. Obtain a p-value matrix {*p*_*ck*_} for every query cell *c* and reference cluster *k*;
  2.4. Apply multiple hypothesis testing correction to the p-values (`p.adj.method=`);
  2.5. Determine the matched cell type for a query cell as the reference cell type that gives the maximum p-value for that query cell, or unassigned (i.e., no matched cell type) if the maximum p-value is below the p-value threshold (`sig.level=`).

### Normalization

The plate-based SMART-seq and droplet-based 10X Genomics protocols are known to have very different read count distributions and detection sensitivities [20]. Thus, normalization is a key step for performing matching across these platforms. In our pipeline, we designed a rescaling and normalization procedure based on the expression value distributions and the signal-to-background-noise patterns observed in the cell type barcode plots.

First, we observed that the gene expression values of the SMART-seq and 10X data had very different dynamic range (Supplementary Figure 1). The marker genes displayed in the cell type barcode were selected by the NS-Forest marker gene selection algorithm that preferentially selects binary expression genes [12], i.e., those genes that are highly expressed in the given cell type and have no/weak expression in other cell types. For the purpose of cross-platform comparison, we designed a gene-wise min-max rescaling step to align the dynamic range of gene expression of both protocols in the range of [0,1]. Let ***x***_*g*_ be a length-*N* vector of the expression value of marker gene *g* across all *N* cells in the dataset. The rescaled expression vector is

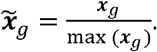

Second, due to the higher sensitivity of the SMART-seq protocol and low detection rate of the 10X protocol for weakly expressed genes, the cell type barcodes displayed weak signals for the genes that are not the marker genes of the given cell type in the SMART-seq data, whereas the cell type barcode of the 10X data displayed zero expression for those genes. For the purpose of cell type matching, the weak expression in the SMART-seq cell type barcodes can be considered a kind of background noise in its we designed the following normalization step. Let X be the rescaled but unnormalized expression sub-expression pattern (Figure 1A). In order to eliminate such background noise in the SMART-seq barcode, we designed the following normalization step. Let 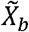 be the rescaled but unnormalized expression submatrix displayed in a cell type barcode *b*. 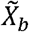 is an *m* × *n*_*b*_ matrix, where *m* is the number of all marker genes, and *n*_*b*_ is the number of cells of cell type *b*. The normalized values are

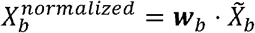

where ***W***_***b***_ is a weighting vector consisting of the row means (or medians) of 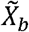. Due to the binaryness of NS-Forest marker genes, ***W***_*b*_ is usually a binary vector with values either close-to-0 or close-to-1. Due to the weighting, the dynamic range of the normalized values may shrink from [0,1]. A final rescaling step is to realign the maximum value of the dynamic range back to 1 sub-matrix-wise, which is:

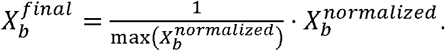

The final expression matrix for the input of the algorithm is the column-concatenation of 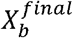 for all *b*’s, where *N* = Σ_*b*_ *n*_*b*_.

The above procedure is implemented in the normalization function `normalization()` in the R package. In the matching use cases presented, the weighting normalization procedure was only applied in the case of cross-platform matching between SMART-seq and 10X protocols. If both the query and reference data are generated using the same platform, the weighting step is not necessary, which can be turned on or off by specifying `norm.by=“mean”, norm.by=“median”`, or `norm.by=NULL` options in the `normalization()` function.

### Cosine distance metric in FR-Match

To make more robust matching, we made another modification in the FR-Match algorithm, which is to calculate the cosine distance that is invariant to scaling as an option for constructing the minimum spanning tree used in the FR test. Let 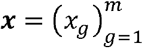 and 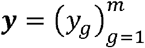 be two cells in the *m*-dimensional feature space of marker genes *g* = 1, ⋯ , *m*. The cosine similarity between the two cells is defined as

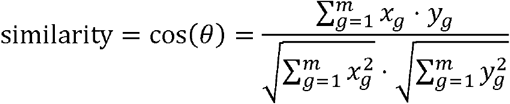

where *θ* is the angle between vectors ***x*** and ***y***. Intuitively, if the angle *θ* is small, then cos (*θ*) is large, which means the two cells ***x*** and ***y*** are more similar to each other as the angle between their representing vectors is small in the multi-dimensional space. If two cells are from different platforms, say ***x*** is SMART-seq data and ***y*** is 10X data, the scale difference between their expression range is normalized by the denominator in the above equation, which is the product of the lengths of the two vectors. Finally, the cosine distance is defined as

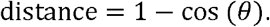

It is suggested to use the scaling-invariant cosine distance for more robust cell type matching across platforms. The option of using cosine distance can be turned on or off by specifying `use.cosine=TRUE` in the `FRmatch()` or `FRmatch_cell2cluster()` functions.

## Data availability

All datasets are publicly available in Allen Brain Map Cell Types Database: RNA-Seq Data (https://portal.brain-map.org/) and NeMO Data Archieve (https://nemoarchive.org/). Specifically, each dataset can be downloaded from the following list.

- Human M1 10X: https://portal.brain-map.org/atlases-and-data/rnaseq/human-m1-10x
- Human M1 SMART-seq: https://portal.brain-map.org/atlases-and-data/rnaseq/human-multiple-cortical-areas-smart-seq
- Mouse MOp single-nucleus RNA-seq: https://assets.nemoarchive.org/dat-ch1nqb7
- Mouse MOp single-cell RNA-seq: https://portal.brain-map.org/atlases-and-data/rnaseq/mouse-whole-cortex-and-hippocampus-10x
- Human MTG SMART-seq: https://portal.brain-map.org/atlases-and-data/rnaseq/human-mtg-smart-seq
- Mouse VISp single-cell RNA-seq and smFISH: https://portal.brain-map.org/atlases-and-data/rnaseq/data-files-2018

Raw count matrices were downloaded and preprocessed by log-transformation of the count per million (CPM) data. Log(CPM) data were the input data of the FR-Match algorithm.

## Code availability

Open source software packages – NS-Forest and FR-Match – are available in GitHub repositories. Reproducible analysis notebooks are also available as tutorials in the software GitHub page. All details can be found in https://jcventerinstitute.github.io/celligrate/.

## Acknowledgements

The work reported in this manuscript was funded by the JCVI Innovation Fund, the Allen Institute for Brain Science, and the U.S. National Institutes of Health (1RF1MH123220). The funding bodies had no role in the design or conclusions of this study.

## Author contributions

YZ and RS conceived the project and prepared the manuscript. YZ and BA conducted the analyses. RA identified and provided the datasets. All authors agreed on the manuscript.

---

**Supplementary Figure 1:**
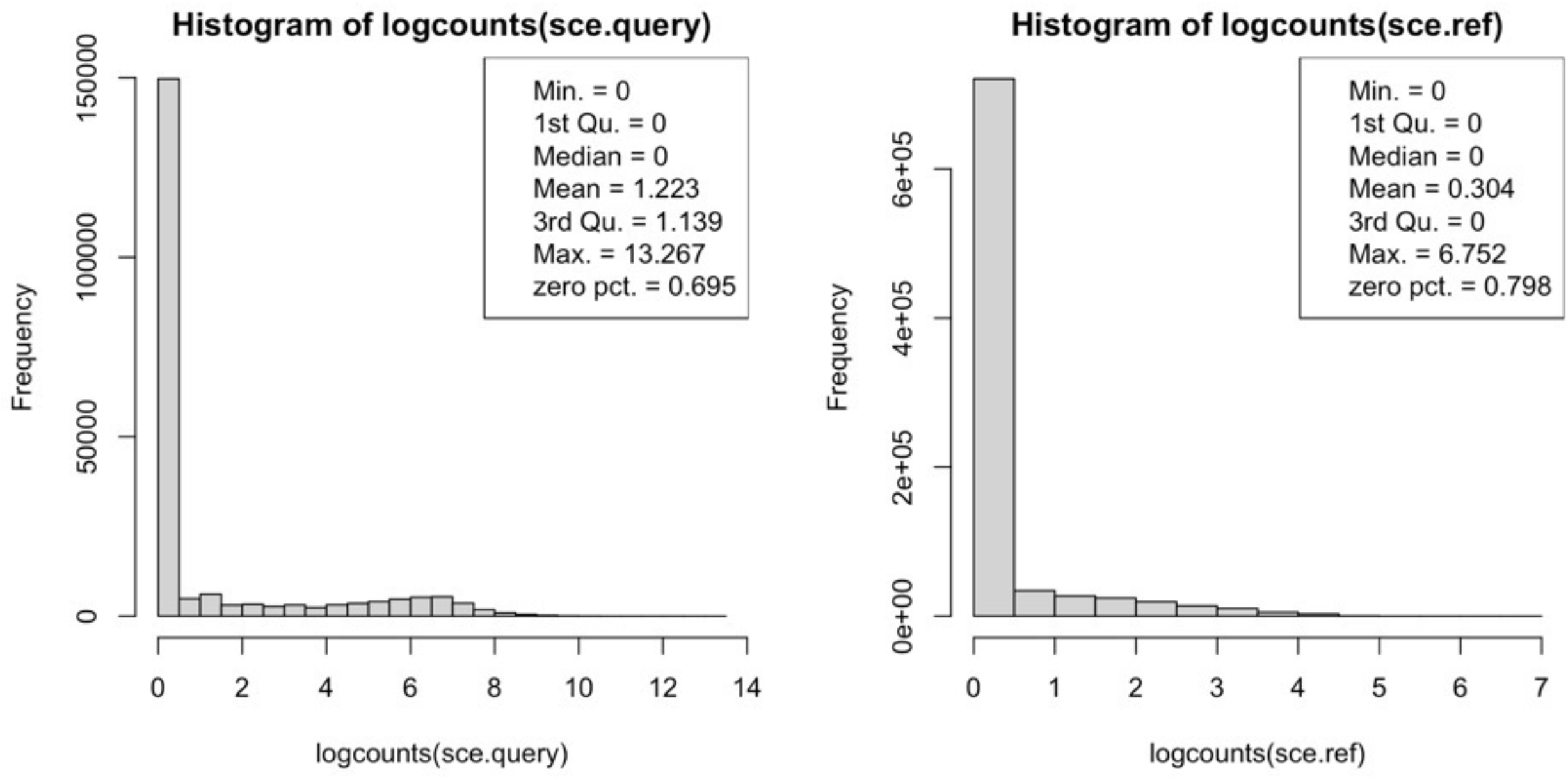
SMART-seq and 10X data distributions. Histograms of the log(CPM) data from the SMART-seq protocol (left) and the 10X protocol (right) are shown. The SMART-seq data form a bimodal distribution, whereas the 10X data form a long-tail right-skewed distribution.

**Supplementary Figure 2:**
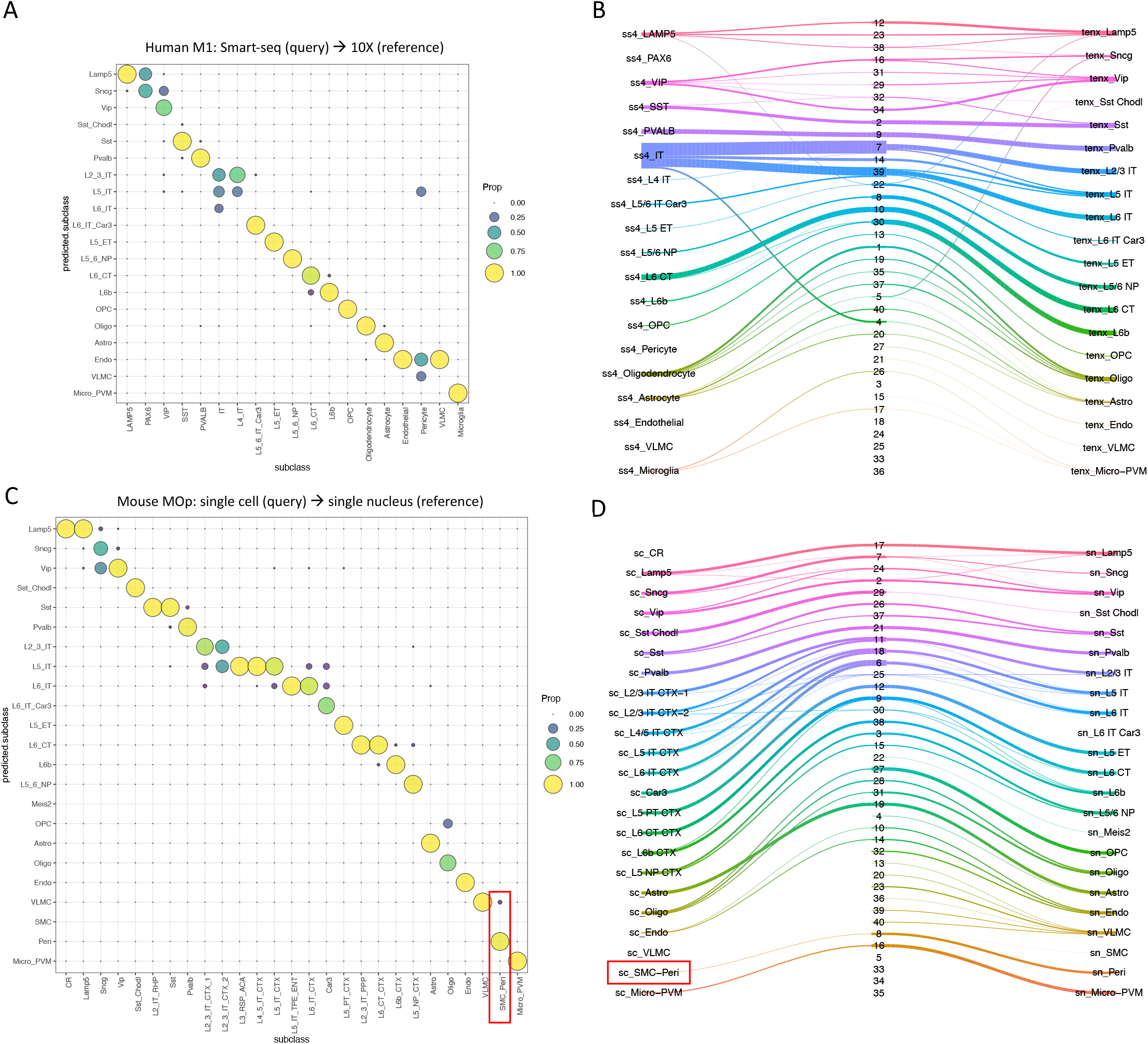
Cell type matching results using Azimuth and Online iNMF. **A.** SMART-seq (query) to 10X (reference) matching results of human M1 subclass cell types using Azimuth. **B.** SMART-seq (query) to 10X (reference) matching results of human M1 subclass cell types using Online iNMF. **C.** ScRNA-seq (query) to snRNA-seq (reference) matching results of mouse MOp subclass cell types using Azimuth. Highlighted in the red box is the potentially under-partitioned query SMC-Peri subclass. **D.** ScRNA-seq (query) to snRNA-seq (reference) matching results of mouse MOp subclass cell types using Online iNMF. Highlighted in the red box is the potentially under-partitioned query SMC-Peri subclass.

**Supplementary Figure 3:**
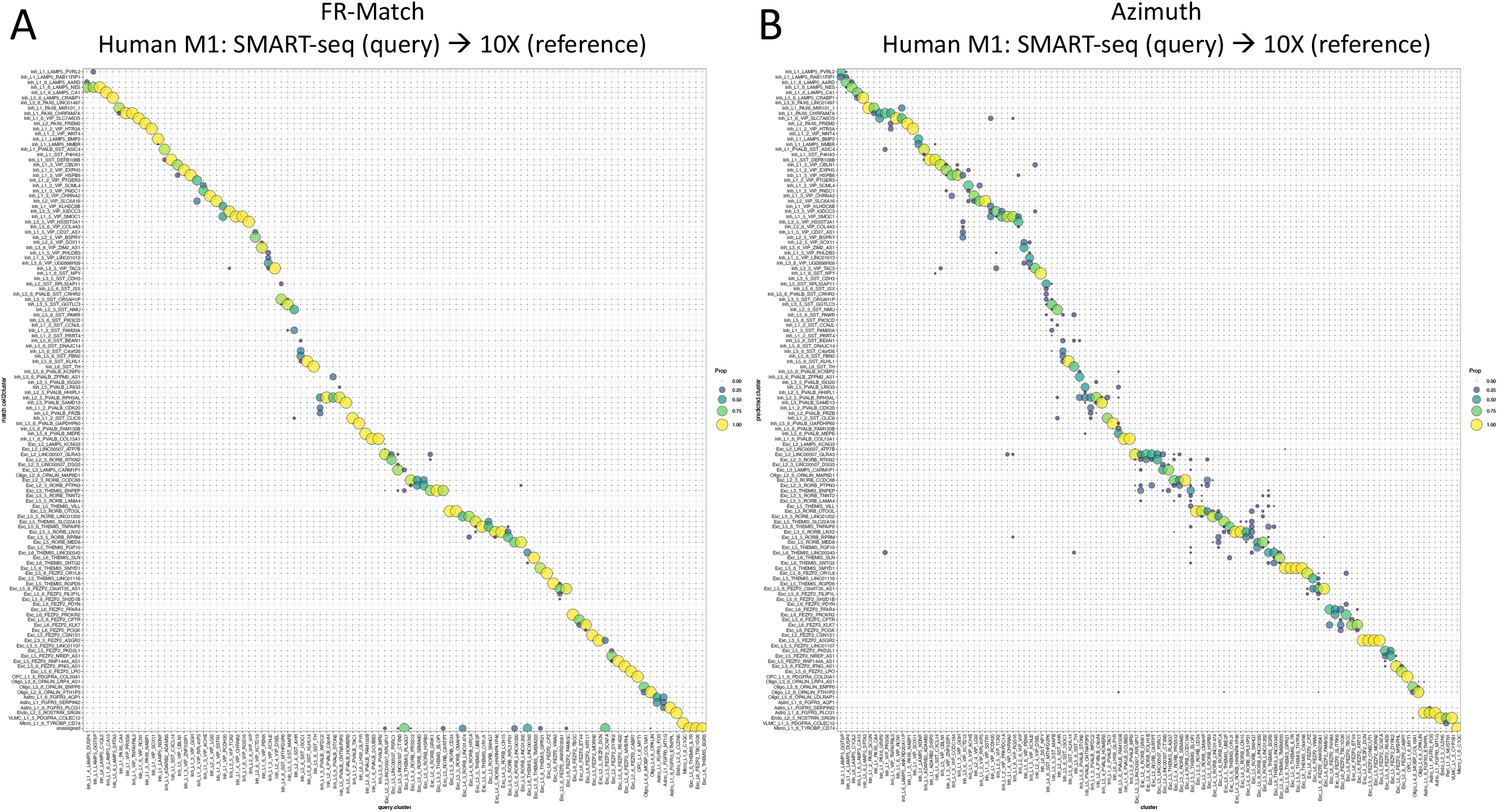
Matching results for matching cell types from SMART-seq (query) to 10X (reference) datasets of human M1 cell types at the most granular cell type resolution using FR-Match (left) or Azimuth (right).

**Supplementary Figure 4:**
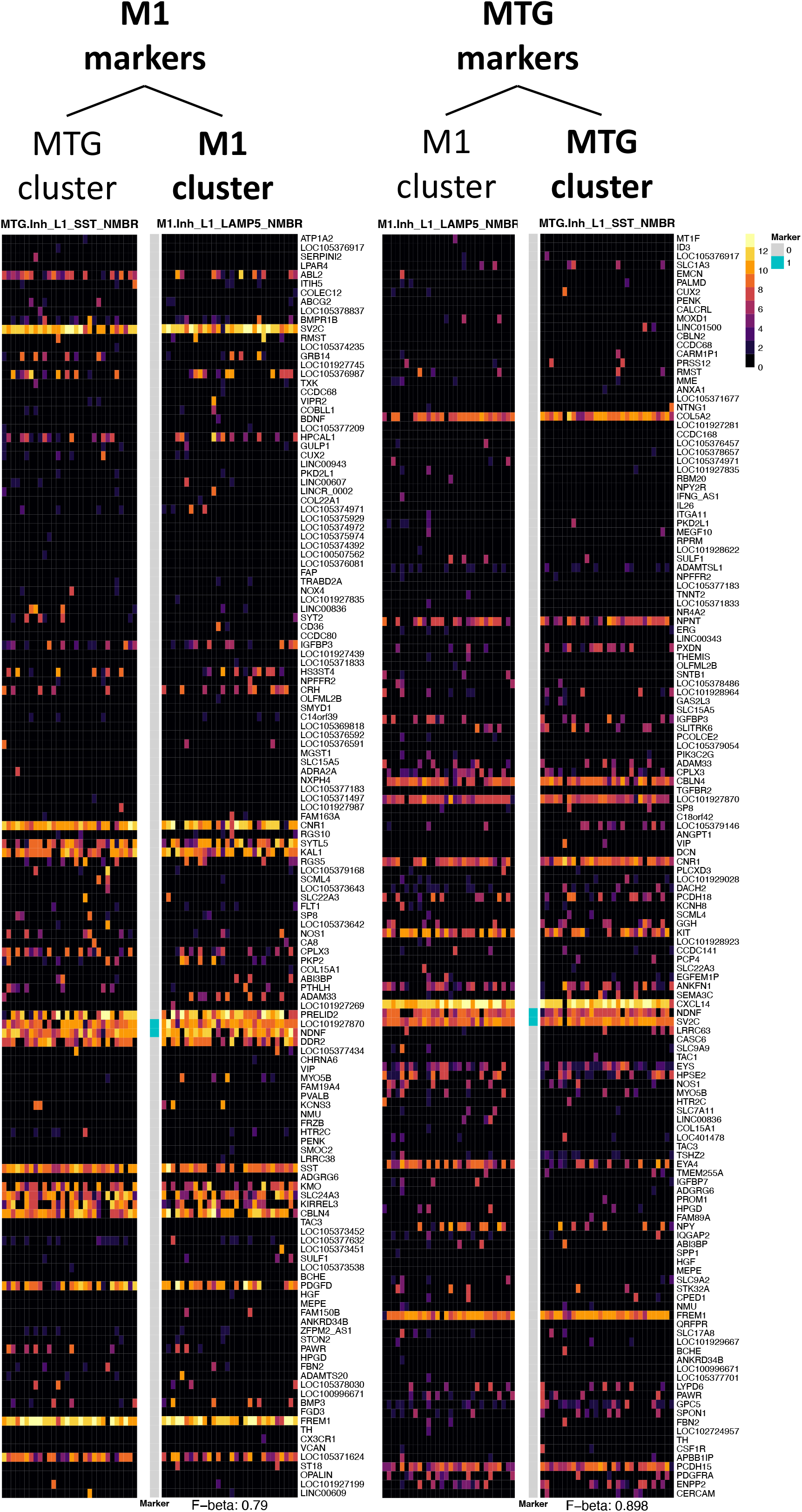
Cell Type Barcode of reciprocal marker genes. Barcode plots for matched cell types (M1.Inh_L1_LAMP5_NMBR and MTG.Inh_L1_SST_NMBR) between M1 and MTG. Matched cell types were identified by two-way FR-Match. Blue vertical bars highlight the marker genes selected for the cell types displayed. The F-beta scores of classification accuracy using the marker gene combinations are listed at the bottom. Left pair are barcodes of the two cell types based on the marker genes derived using the M1 dataset. Right pair are barcodes of the two cell types based on the marker genes derived using the MTG dataset. Both pairs show very similar expression patterns of the barcodes within each pair on the reciprocal sets of marker genes, supporting the close similarity of the matched cell types between the different brain regions.

